# Inhibition of Lipopolysaccharide *E. coli-*induced acute lung injury by extracted *Antidesma bunius* (L.) Spreng fruits as compared to Fluticasone Propionate, a corticosteroid

**DOI:** 10.1101/2021.04.08.438930

**Authors:** Maria Nilda M. Muñoz, Jennifer Lucero, Kimberly Stacy Hope Benzon, Jerica Isabel L. Reyes, Charina de Silva, Reuel C. Delicana, Urdujah A. Tejada, Swapan Chatterjee, Kozo Watanabe

## Abstract

The hallmark of Acute Lung Injury/Acute Respiratory Distress Syndrome (ALI/ARDS) is inflammation-induced alveolar-vascular barrier destruction and neutrophilic infiltration that leads to the formation of cytokines and oxygen radicals. The objective of the study is to investigate the protective and toxicological effects of *Antidesma bunius* (L.) Spreng [Bignay] in murine model of Lipopolysaccharide *E. coli* (LPS)-induced ALI and compared with Fluticasone Propionate (FP), a synthetic corticosteroid. We showed that extracted Bignay fruits have high amount of phenols, steroids and flavonoids but insignificant amount of heavy metals and aflatoxins. BALB/c mice of either sex were divided into 4 groups in the ALI mouse model; Group 1: vehicle control; Group 2: LPS alone; Group 3: Bignay + LPS; and Group 4: FP + LPS. Bignay and FP were administered via intraperitoneal injection while LPS was given intra-tracheally. Biomarkers of ALI such as total lung inflammatory cell count, total lung protein content, lung edema and interleukin-6 (IL-6) secretion were measured 24 hrs after vehicle control or LPS treatment. Compared to vehicle controls, LPS caused significant increased in all measured biomarkers of ALI in samples collected from bronchoalveolar lavage fluid and were significantly attenuated by Bignay fruit extract or FP. Pulmonary vascular leakage caused by LPS was also evaluated after injection with Evans blue dye, an indication of lung injury. Extracted Bignay fruits or FP when given to mice 2 hrs after LPS administration substantially decreased the pulmonary vascular leak. Our findings are the first evidence demonstrating the preventive and non-toxic effects of extracted Bignay fruits in a murine model of LPS-induced ALI. The results could be attributed to the presence of active secondary metabolites such as flavonoids, phenols and steroids. It is also evident that extracted Bignay fruits are as effective as FP, well-established steroid, in blocking the biomarkers of ALI caused by LPS.

## Introduction

Acute lung injury/acute respiratory distress syndrome (ALI/ARDS) is a life-threatening disease characterized by inadequate oxygenation, pulmonary infiltration, and observation of onset in critically ill patients [1–4]. This can arise from a plethora of insults, either direct or indirect, to the lung [1–3]. Increased epithelial permeability results from the insult, leading to alveolar flooding of a protein-rich edema fluid, increased expression of cytokines and chemokines and upregulation of adhesion molecules [2,4,5]. Acute respiratory failure and usually catastrophic disease such as ALI/ARDS, lead to the resulting loss of gas exchange, requiring assistance for ventilatory and critical care. ALI/ARDS has a high mortality rate and there are few successful therapeutic modalities to fight this disease [1–4].

There has been a growing interest in medicine derived from natural products, and ongoing studies are actively investigating *in vivo* and *in vitro* multiple curative plants for various inflammatory diseases [9, 10]. To date, in many rural areas and low-income communities, the cost of synthetic medication is growing and becoming unavailable. Because of these issues, there are mounting concerns about developing herbal medicine around the world. The use of herbal medicine is currently expected to be more practical due to its affordability, availability, and, most significantly, less harmful to humans. Previous studies have shown that various parts of medicinal plants are consumed by ingestion, decoction and topically applied to infected parts of the body [9,11,12]. Nevertheless, there is still a lack of consistent and full knowledge on the composition of plant extracts used in herbal medicine, as well as the occasional toxicity of adulterants and/or environmental contaminants [12–14].

In Bignay extracts, both heavy metals [15, 16] and aflatoxins [17, 18] were assessed because these toxins are extremely stable prior to experimental procedures under most storage, handling and dispensation conditions. In plants that grow in the soil, lead (Pb) and cadmium (Cd) are among the heavy metals since they are normal components of the earth’s crust (Agency for Toxic Substances and Disease Registry, 2011). In popular foods, we consume more heavy metals every day, but if added at elevated levels, it can cause toxicity.

Aflatoxins are naturally occurring compounds that are produced from the molds *Aspergillus flavus* and *Aspergillus parasiticus* [19, 20]. These are also produced in soil when cultivated in a wide range of agricultural products [19, 20]. An aflatoxin acceptance limit of 20 parts per billion (ppb) has been set by the US Food and Drug Administration for meats, including fruits, and for most feedstuffs and ingredients. On the other hand, the European Union has reached the level of tolerance of aflatoxin B1 and complete aflatoxin in nuts, dried fruits and cereals for human consumption at 2 and 4 ppb, respectively. Due to the proposed restrictions, the presence of aflatoxin in agricultural products is not only a significant food safety problem, but also has important cost-effective implications for the agricultural and pharmaceutical industries.

The Philippines is known for its abundance of flora and that herbal medicine had long been used even before the advent of synthesized medicine. In herbal medicine, the leaves and stems are usually involved; fruits, however, have medicinal uses as well. For example, berries such as *Antidesma bunius* (L.) Spreng [Bignay], *Synsepalum dulcificum* (Miracle Berry), *Basella Alba linn* (Alugbati), and *Morus alba L.* (Mulberry) are grown in the Philippines and is noted for its high levels of phenols and flavonoids due to its anti-oxidant activity [21–25]. Phytochemical studies of berries, saponins and phytosterols, have been reported contributing to the anti-inflammatory activities of extracted fruits [26–29]. However, the anti-inflammatory effect of berries on biomarkers of ALI/ARDS, in particular *Antidesma bunius* (L.) Spreng fruits, has yet to be investigated.

In the Philippines and other countries such as southern China, Vietnam, India’s lower Himalayas, Thailand, Indonesia, Singapore, New Guinea, Cuba and Florida, the *Antidesma bunius* (L.) Spreng fruit is commonly called “Bignay or Bugnay” and belongs to the family of Phyllanthaceae. For the prevention of diabetes, urinary tract infection and hypertension [31–33], Bignay and other fruits are normally boiled or eaten raw by indigenous Filipinos. While important active components of Bignay [9,21,26,27,31] have been identified, little is known about the efficacy of Bignay in the *in vivo* system, particularly as a result of its use to improve lung disorders such as asthma and ALI/ARDS.

The protective and toxicological effects of Bignay fruit extracts in LPS-induced ALI and possible mechanisms involved were investigated in this study. We chose this model because LPS produces a relatively mild form of ALI without causing death [35, 36]. To our knowledge, there is no scientific evidence concerning the protective and toxicological effects of Bignay in ALI/ARDS on either humans or animals. Further exploration of Bignay should establish the pathological role of the inadequately characterized Bignay fruit in lung disorders and other inflammatory diseases.

## Materials and methods

### Chemicals/Reagents/Plant sample

Lipopolysaccharide *E. coli* (LPS) and other chemicals were obtained from Sigma-Aldrich Chemical Co. (St. Louis, MO., USA) unless otherwise specified. Precautionary measures in handling LPS were followed as outlined in the Material Safety Data Sheet. Mouse ELISA Interleukin (IL)-6 kit was purchased from Invitrogen, Philippines. Fluticasone Propionate (GlaxoSmithKline) was a gift from Dr. Teresita de Guia, Philippine Foundation for Lung Health Research Development, Philippine Heart Center, Quezon City.

*Antidesma bunius* (L.) Spreng (Bignay) fruits were harvested from Cagayan Valley farm, Region 2, Philippines during the month of May-July (Fig 1). A voucher sample was sent for authentication to the National Museum of the Philippines. After collection, Bignay fruits were washed, oven-dried and pulverized. Bignay fruits were extracted in 70% of Laboratory grade ethyl alcohol and were stirred at 50°C for 4 hrs. The mixture was filtered via gravimetric filtration using Whatman filter paper No. 1 (SigmaAldrich Chemical Co.) and the filtrate was concentrated to remove any residual alcohol using a GeneVac (SP Scientific, Stone Ridge NY). The device was set at 40°C until syrupy in appearance was achieved. The residue was stored at −80°C and resuspended in distilled water prior to its use.

**Fig 1.**
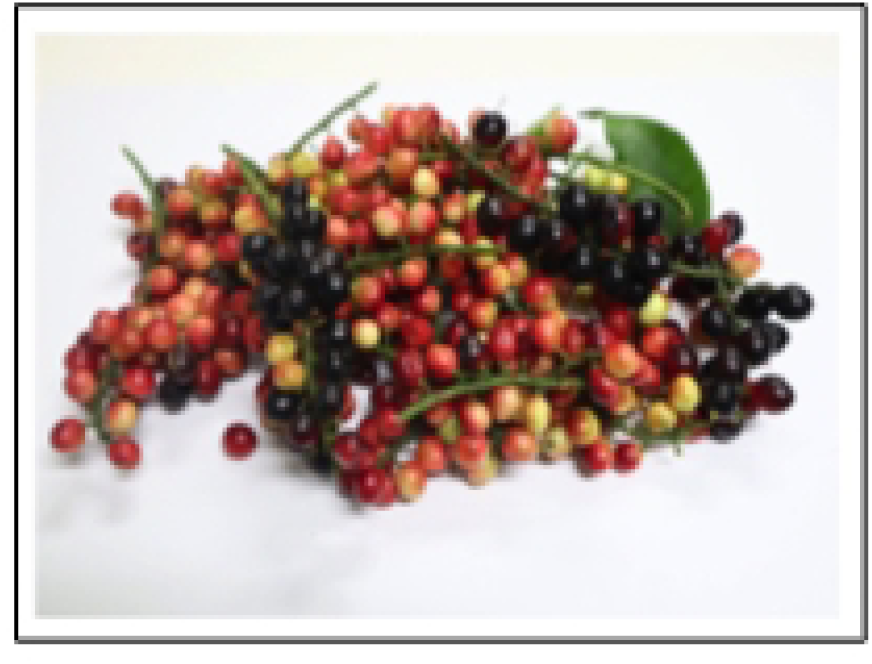
*Antidesma bunius* (L.) Spreng (Bignay). The seasonal fruits belong to the family Phyllanthaceae. It is a native of the Philippines, commonly known as “Bignay or Bugnay”, and the fruiting season is between May and July. In grapelike clusters, the round or ovoid fruit is borne and it matures unevenly. Unripe Bignay fruits, reddish orange in color, were used in this study because preliminary data showed that the presence of secondary metabolites was greater than that of matured ripe Bignay.

### Quantitation of Contaminants and secondary metabolites

#### Measurement of Heavy Metals

The fresh Bignay fruits were oven dried until the moisture content is equivalent to13%. Dried Bignay fruits samples were first digested using wet digestion method [37]. Briefly, 0.2 g of dried samples were taken in 100 mL flask and immediately, 4 mL of 0.02M HNO_3_ acid was added. The mixture was allowed to stand for 4 hrs, heated over water bath till red fumes coming from flask ceases. After cooling, 4 mL perchloric acid was added to the flask and heated again over water bath till small amount was left; subjected to filtration using Whatman filter paper #42 (SigmaAldrich Chemical Co.). The filtrate was injected to Flame Atomic Absorption Spectrophotometer (Shimadzu AA-7000) for determination of Lead (Pb), and Cadmium (Cd) in prepared samples.

#### Measurement of Aflatoxins

The total content of aflatoxin and aflatoxin B1 were quantified using AgraQuant^®^ Total Aflatoxin Test Kits from Romer Labs, Singapore to ensure that *Antidesma bunius* (L.) Spreng [Bignay] fruits are free from traces of toxic substances. These kits are kits for enzyme-linked immunosorbent assay (ELISA) and the protocols for the study were followed according to instructions by manufacturer. Using the BMG LABTECH Spectrostar Nano Microplate Reader (BMG Lab Tech, Germany) set at 450 nm, the absorbance of standards and samples was measured. The Bignay samples were used for triplicate measurements of the overall alfatoxin and aflatoxin B1 content. Data are expressed as the mean values in parts per billion (ppb).

#### Determination of Total Phenolics

Total phenolic content was determined by Folin-Ciocalteau’s phenol reagent and 2% Na_2_CO_3_ [38]. The external calibration was performed using different concentrations of Gallic Acid (GA) *i.e*., 0.00, 25 µL, 50 µL, 100 µL, 250 µL, and 400 µL. The reaction mixtures were performed in triplicate, and phenolic content was measured using Spectrostar Nano UV/Vis Spectrophotometer (BMG Lab Tech, Germany) set at 765 nm. Results were expressed in mean ± standard error of mean (SEM) as mg GA equivalent per gram sample using GA calibration curve.

#### Determination of Total Flavonoids

The total flavonoid content of Bignay was measure by aluminum chloride (AlCl_3_) method [39]. The mixture of Bignay (1 mL sample + 4 mL water) was placed in 50 mL volumetric flask. A 0.3 mL 5% NaNO_2_ was added to the mixture and was kept in dark place for 6 min; thereafter, 0.3 mL 10% AlCl_3_ was added. Five min later, 5% NaOH was added to complete the reaction. Absorbance of sample against blank was read at 510 nm using Spectrophotometer (BMG Lab Tech, Germany). Quercitin standard was used for the calibration curve, generated using five point concentrations of 25 µL, 125 µL, 250 µL, 350 µL, and 500 µL in a multiplate. Values were expressed in mean ± standard error of mean (SEM) in terms of flavonoid content (quercitin equivalent per g of dry weight).

#### Determination of Total Steroids

Estimation of total amount of steroids was performed by Liebermann Burchard reaction [40].

Standard calibration levels were prepared using cholesterol as recommended in the manufacturer’s kit protocol. All these reaction test tubes (standards) were gently vortexed for 10 secs and put in a water bath (37°C) for 10 min. The solution was allowed to cool down to room temperature for 10 mins after incubation. A 300 μL of Bignay extracts were pipetted and then combined with 2.50 mL of the cholesterol color developer. For 10 secs, the resulting mixture was vortexed and the reaction tubes were incubated for 10 mins in a 37°C water bath. The test tubes were allowed to cool down to room temperature prior to absorbance reading. The same powder as Bignay was prepared for Lagundi powder, which acts as the positive control for this assay. The spectrophotometer was set at 620 nm, which was used as a blank solution for distilled water. Readings of all standards and samples were registered and the triplicate and two separate experiments were performed. Data are expressed as mean +/-standard error of mean (SEM).

### Experimental Animals and Biomarkers of ALI

#### Preparation of Animals

BALB/c mice (8-10 weeks, ∼20-25 g in weight) of either sex were purchased from Research & Development, St. Luke’s Medical Center and College of Veterinary Medicine, Animal Research Facility at Cagayan State University, Philippines. The reported studies were consistent with the concepts outlined by the guidelines of the Institutional Animal Care and Use Committee (IACUC) and Animal Research: Reporting in Vivo Experiments (ARRIVE) in biomedical research [41]. A permit to ensure the proper treatment and use of animals for this study was provided by the Philippine Bureau of Animal Industry.

Experimental design

BALB/c mice of either sex were divided into 4 groups:

Group 1 (n=5 mice): vehicle control [no LPS, no Bignay, no Fluticasone Propionate (FP)]

Group 2 (n=5 mice): 10 mg.kg^-1^ LPS alone

Group 3 (n=5 mice): 1000 mg.kg^-1^ Bignay *plus* 10 mg.kg^-1^ LPS Group 4 (n=5 mice): 10^-7^ M FP *plus* 10 mg.kg^-1^ LPS

The 1000 mg.kg^-1^ dose of Bignay was chosen because previous reports have shown that 1000 mg.kg^-1^ Bignay had no effect in different organs after acute oral toxicity study (OECD 423) in mice *in vivo* (Japan Association for Laboratory Animal Science Conference, May 2017:Fukushima, Japan). Preliminary studies have demonstrated that no animals died at the highest dose (2000 mg.kg^-1^), thus suggesting that this is an indicative of LD_50_ are greater than 2000 mg.kg^-1^. The 10 mg.kg^-1^ dose of LPS was used which was shown to cause ALI and neutrophilic inflammation in mice *in vivo* [35]. Fluticasone Propionate (10^-7^ M FP), a class of drug known as corticosteroids, was used as reference drug for all subsequent *in vivo* experiments [42–45]. This dosage is based on previous studies that demonstrated reduction of secreted airway mediators [35].

The vehicle control (distilled water) or 10 mg.kg^-1^ LPS was instilled intratracheally into anesthetized mice. Twenty-four hrs later, mice were anesthetized with combination of dexmedetomidine (Precedex; 100 µg.mL^-1^) and ketamine (100 mg.mL^-1^) via intraperitoneal injection. A total of 1 mL x 3 PBS was slowly infused into the lung and bronchoalveolar lavage fluid (BALF) was collected by gentle aspiration. After centrifugation (12,000*g* x 5 min), the supernatant was saved and was stored for later determination of secreted IL-6 and total lung protein content. The cell pellet was resuspended in PBS, and the totl lung inflammatory cells were counted manually using Neubauer hemocytometer (Fisher Scientific USA).

#### Measurement of Total Lung Protein

The total lung protein in BALF was measured using the Bradford Reagent Kit [SigmaAldrich Chemical (Chemline Scientific, Philippines)]. In the PBS buffer, bovine serum albumin used as standard (62.5–1,400 μg.mL-1), was prepared. In the 96-microplate well, 100 mL supernatant or cell lysate samples and 100 μL of protein standard were separately added; blank wells have PBS buffer alone. 100 μL of Bradford Reagent was applied to each well and incubated for 5 min. The samples were measured at a 595 nm absorption rate. By comparing the net absorbance values against the standard curve, the protein concentration of the unknown samples was calculated.

#### Quantitation of secreted IL-6

Supernatant obtained from the BALF was used to measure the endogenous release of IL-6 using a commercially available Murine IL-6 ELISA kit according to the manufacturer’s instructions (ThermoFisher, InVItrogen). The absorbance was read at 450 nm and secreted IL-6 was analyzed using Gen5 data analysis software (BioTek Instruments, Inc.). The minimum detectable dose of IL-6 is <2 pg.mL^-1^. All data are expressed in pg.mL^-1^.

#### Determination of lung wet:dry ratio

In separate experiments, treated mice as above were euthanized, chest was opened and lungs were excised. Each lung was blotted off, weigh and placed in an oven set at 50°C for 48 hours to achieve dry weight. The wet:dry ratio was calculated to assess lung edema formation caused by LPS in the presence or absence of either vehicle control, Bignay or FP.

In another set of experiments, treated lungs were fixed by gentle injection of 4% buffered paraformaldehyde into the tracheal cannula at a pressure of 20 cm H_2_O. The excised lung was immersed in 4% paraformaldehyde solution for 24 hrs. The inferior lobe of right lungs were sectioned sagittally, embedded in paraffin, cut into 5 µm sections, and stained with hematoxylin and eosin (H&E) for morphological examination.

#### Evaluation of acute pulmonary vascular leakage

To examine whether LPS causes pulmonary vascular leakage, Evans blue dye was injected into tail vein in all treated mice. Evans blue is an azo alkaline stain that has a high affinity for serum albumin [46]. This dye is used as an *in vivo* marker to examine the LPS-induced vascular permeability of lung damage. All animals received Evans blue dye but lungs were not infused with PBS (no BAL).

Experimental interventions: Mice were selected randomly.

Group 5 (n=5 mice): vehicle control [no LPS, no Bignay, no Fluticasone Propionate (FP)]

Group 6 (n=5 mice): 10 mg.kg^-1^ LPS alone

Group 7 (n=5 mice): 1000 mg.kg^-1^ Bignay *plus* 10 mg.kg^-1^ LPS Group 8 (n=5 mice): 10^-7^ M FP *plus* 10 mg.kg^-1^ LPS

At the end of experimental period, mice were sacrificed 24 hrs after treatment. Lung samples were collected for actual visible coloration analysis.

#### Statistical Analysis

Measured data were presented as mean ± standard error of the mean (SEM). Different treatments were analyzed by Student’s *t-test* for analysis of significant differences between two groups. The use of ANOVA with Tukey test was used to test for population mean differences; mean of all animals treated with Bignay, mean of all animals treated with FP, which is considered to be the best available method in cases where confidence intervals are required. In comparison of three or more groups, Tukey method was used as *post hoc* test regarding p<0.05 as significant after analysis of variance (ANOVA). GraphPad Prism Software 9.0 (GraphPad, San Diego, CA) was used for all analyses.

## Results

### Quantitative analysis of heavy metals

The amounts of Lead (Pb) and Cadmium (Cd) were both undetected (Table 1), measured by the Philippine Accredited Institute of Pure and Applied Chemistry using AAS. Our results indicated that in processed and stored samples, the Bignay fruit extracts have no trace of heavy metals. The permissive limit and limit of detection are shown in Table 1.

**Table 1.**
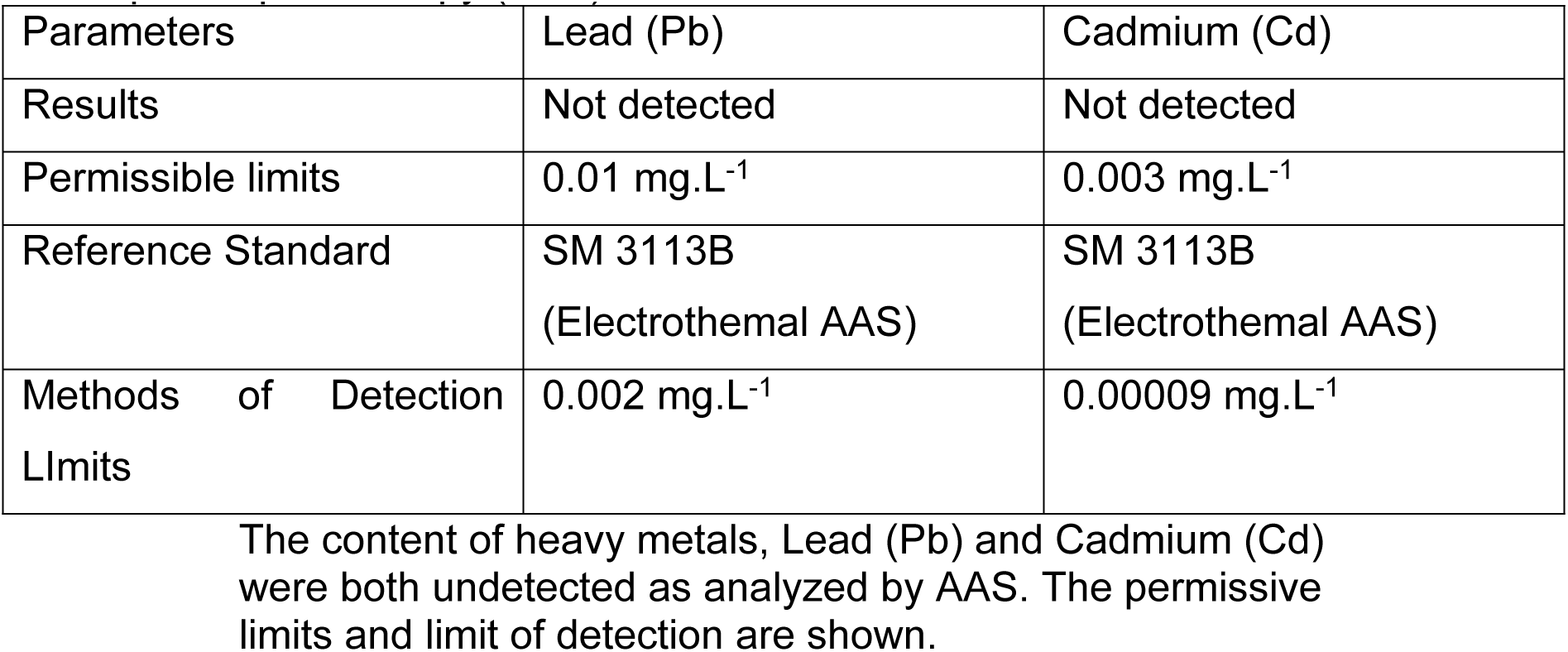
Analysis of Heavy Metals from Dried Bignay Samples by Atomic Absorption Spectroscopy (AAS)

| Parameters | Lead (Pb) | Cadmium (Cd) |
| --- | --- | --- |
| Results | Not detected | Not detected |
| Permissible limits | 0.01 mg.L<sup>-1</sup> | 0.003 mg.L<sup>-1</sup> |
| Reference Standard | SM 3113B<br>(Electrothermal AAS) | SM 3113B<br>(Electrothermal AAS) |
| Methods of Detection Limits | 0.002 mg.L<sup>-1</sup> | 0.00009 mg.L<sup>-1</sup> |

The content of heavy metals, Lead (Pb) and Cadmium (Cd) were both undetected as analyzed by AAS. The permissive limits and limit of detection are shown.

### Analysis of Aflatoxins

Aflatoxin exposure through food can result in serious health complications and consequences. We observed that the content of total aflatoxins and aflatoxins are below the stipulated limit as reported in the Guidelines of the Registration of Herbal Medicines of Department of Health.

### Quantitation of polyphenols, phytosterols and flavonoids

The total phenolic quantity of extracted Bignay fruit was 20.4 ± 0.2 mg GA equivalent/gram (p<0.01 vs. control of the vehicle). The reference medicinal plant, Lagundi yielded 8.0 ± 2.0 mg GA equivalent/g extract, which was ∼ 60 percent lower than Bignay (p<0.05 vs. Bignay). Lagundi was used because it is known to have significant active secondary metabolites and anti-inflammatory property [47].

The content of total phytosterol in Bignay fruit extracts was 33.0 ± 7.0 mg cholesterol equivalent/gram extract . The phytosterol content of Bignay was also found to be ∼3-fold higher than that of Lagundi, detecting just 10.1 ± 2.0 mg cholesterol equivalent/g extract (p<0.05; Bignay vs. Lagundi).

Bignay’s total flavonoid level was 12.2 ± 4.0 mg quercetin equivalent/g extract. The normal reference Lagundi flavonoid was comparable to the Bignay ethanolic extract (p=NS).

### Indices of LPS-induced ALI: effect of Bignay and FP

#### Total Lung Cell Number

The total number of lung cells from vehicle control was 105,000 ± 70,000 and increased to 3,073,000 ± 510,420 cells (p<0.0003 vs. vehicle control after administration of LPS alone (Fig 2). The overall lung cell count was attenuated by treatment with 1000 mg.kg-1 extracted Bignay fruits to 1,230,170 ± 292,700 cells (p<0.001 vs LPS alone). The protective effect of Bignay+LPS in inhibiting the total lung cell number caused by LPS was less efficient when compared to FP+LPS (p<0.002). ALI induced by LPS was blocked to near baseline with FP+LPS (p<0.001 vs LPS alone, no Bignay).

**Fig 2.**
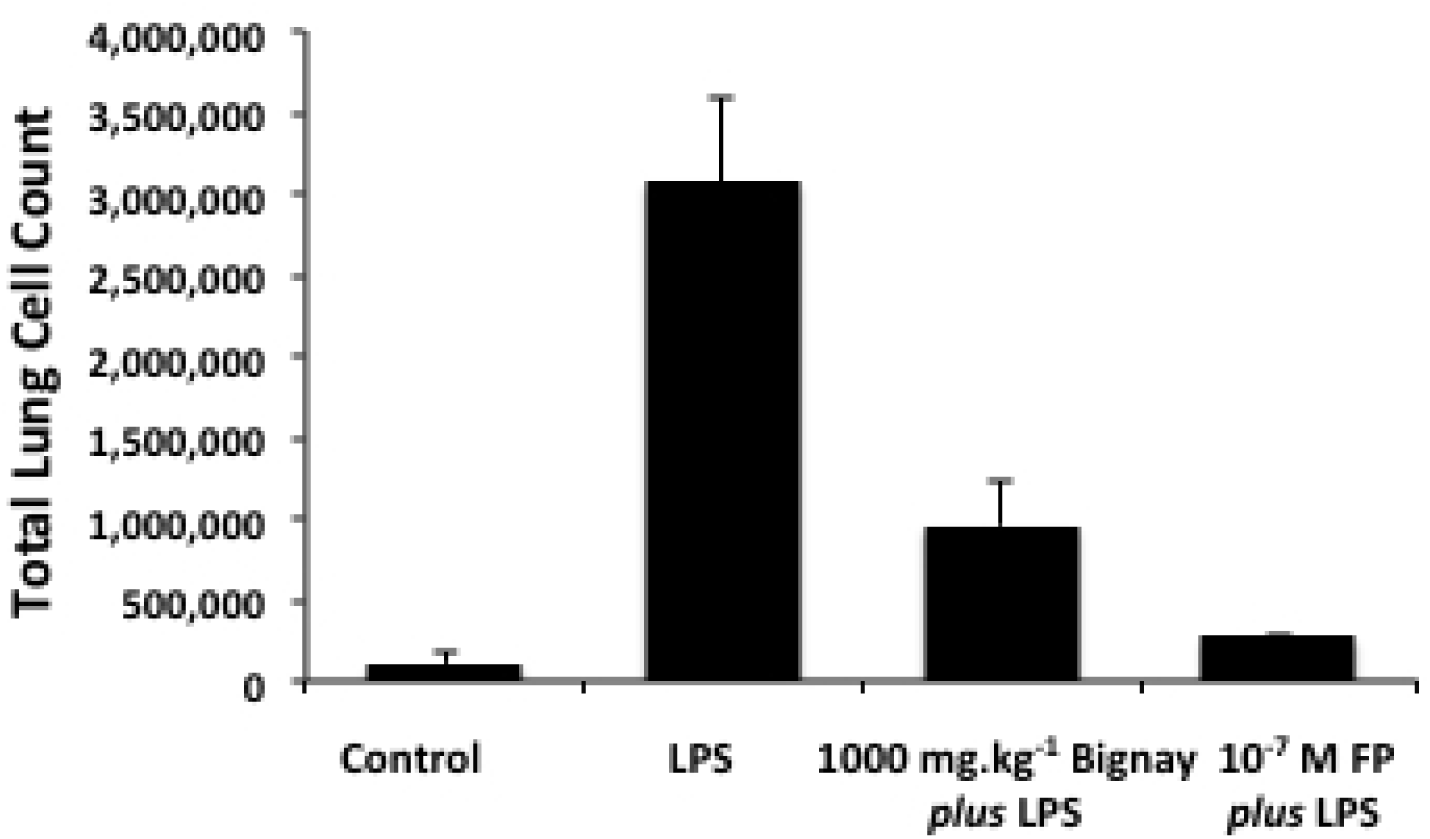
Total cell count obtained from bronchoalveolar lavage fluid (BALF; n=5 mice/group of either sex). Administration of LPS caused increased in total lung cell count and was >70% blocked with extracted *Antidesma bunius* (L.) Spreng [Bignay]. Fluticasone Propionate (FP) was used as reference drug and known to block the cell migration to the site of inflammation. PBS is the vehicle control used in this study. Total cell count was determined manually by using the hemocytometer. Where significant differences were identified, Tukey’s test was used to further examine the differences. p<0.0003, LPS alone vs. vehicle control; p<0.001, B+LPS vs. vehicle control; p<0.01, FP+LPS vs. vehicle control; p<0.001, LPS alone vs. B+LPS; p<0.001, LPS alone vs. FP+LPS; p<002, B+LPS vs FP+LPS.

#### Total Lung Protein

The total lung protein was measured by Bradford assay (Fig 3). There was an increased in total lung protein in mice receiving LPS alone compared to vehicle treated mice (p<0.01, LPS vs vehicle control). An ∼50% reduction of total lung protein caused by LPS stimulation was demonstrated from animals receiving Bignay extract (p<0.05 vs LPS alone, no Bignay) while FP elicited <30% inhibition from stimulated value of LPS. Importantly, we observed that Bignay is apparently more effective than FP in inhibiting the total lung protein.

**Fig 3.**
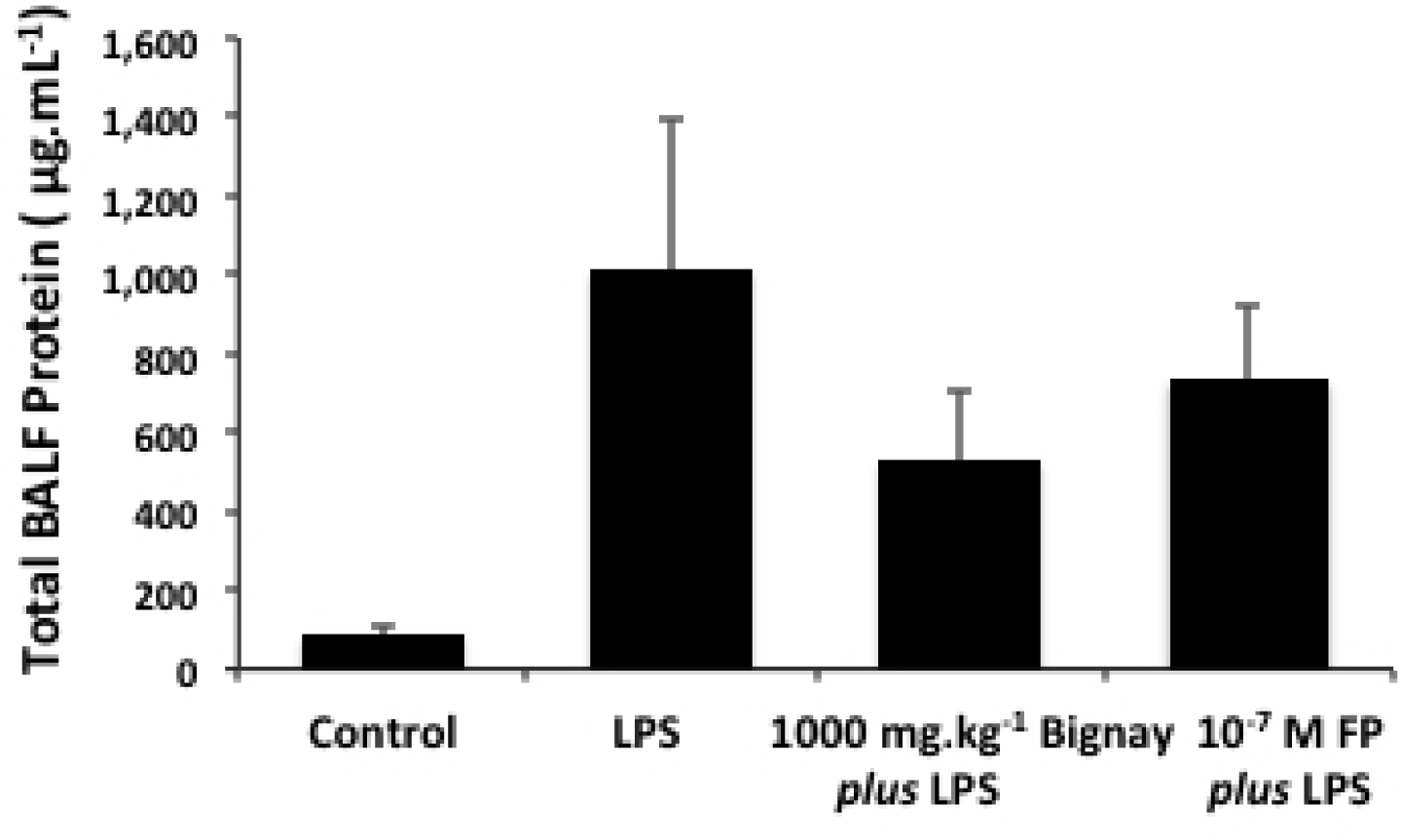
Total Lung Protein Content. LPS-induced acute lung injury increased total lung protein (µg.mL^-1^) in 5 experimental mice. LPS caused a marked increase in total lung protein obtained from BALF. Bignay and Fluticasone Propionate (FP) inhibited minimally the protein content obtained from BALF when compared with LPS alone. Where significant differences were identified, Tukey’s test was used to further examine the differences. p<0.01, LPS alone vs. vehicle control; p<0.001, B+LPS vs. vehicle control; p<0.01, FP+LPS vs. vehicle control; p<0.05, LPS alone vs. B+LPS; p<0.05, LPS alone vs. FP+LPS; p=NS, B+LPS vs FP+LPS.

#### Secretion of IL-6

IL-6 is one of the biomarkers of ALI [5,7,8]. Administration of LPS caused substantial secretion of IL-6 as compared to baseline value (Fig 4). Extracted Bignay fruits inhibited the secreted IL-6 by ∼55% from LPS and was reproducible for all treated mice. Similarly, we observed that the inhibitory effect of synthetic corticosteroid, FP, was as effective as Bignay in blocking the release of IL-6. Our data indicate that Bignay could mimic the protective effect of FP in reduction of LPS-induced ALI.

**Fig 4.**
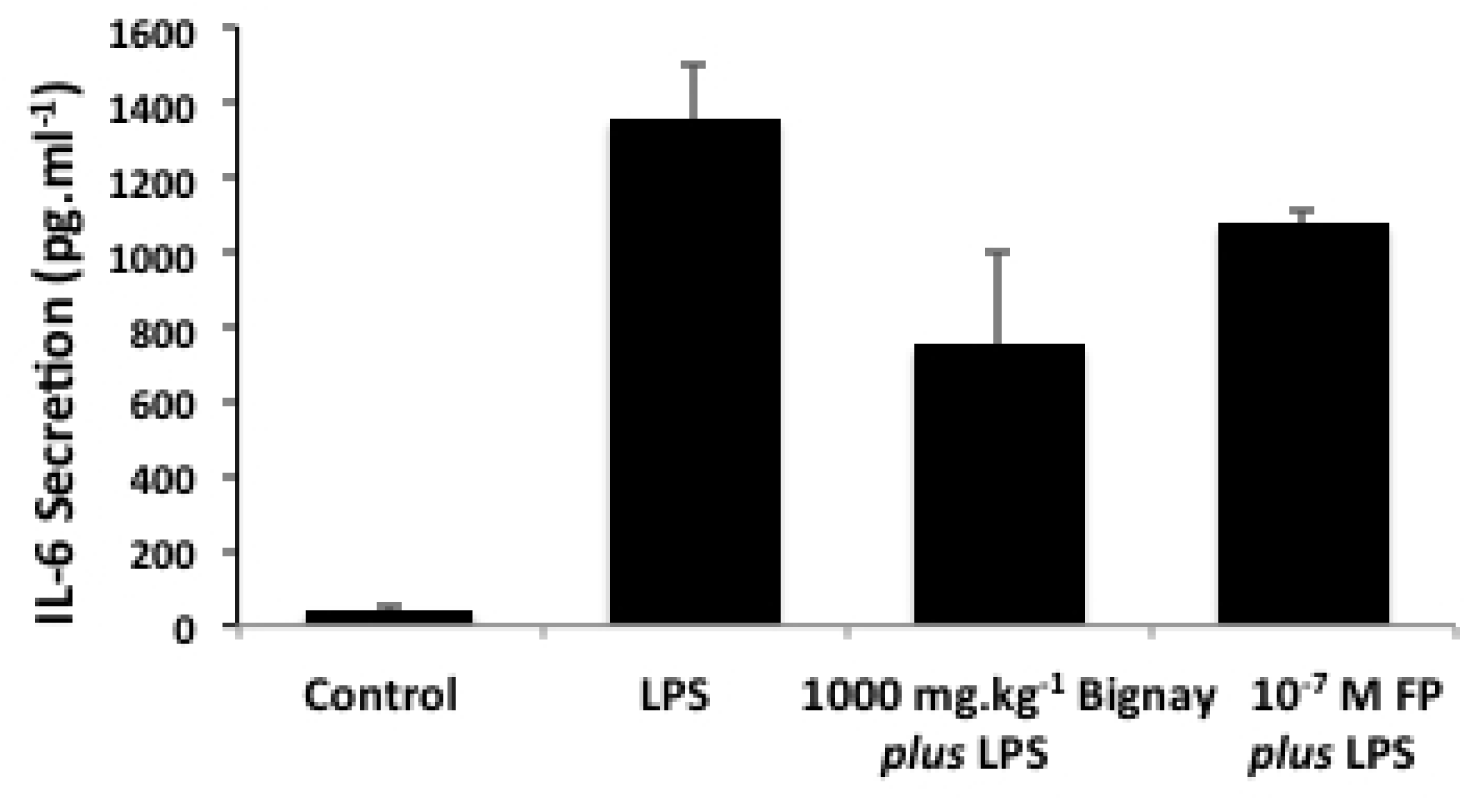
Blockade of LPS-induced endogenous secreted IL-6 by *Antidesma bunius* (L.) Spreng [Bignay]. Administration of LPS caused increased secretion of IL-6 compared to vehicle control treated mice. Treatment with 1000 mg.kg^-1^ Bignay and 10^-7^ M Fluticasone Propionate (FP) attenuated equally the secreted IL-6 caused by LPS stimulation. Concentration IL-6 in the BALF was measured using the IL-6 Mouse ELISA kit. Measurements are mean ± SEM, and expressed as picogram per milliliter (pg.mL^-1^). Where significant differences were identified, Tukey’s test was used to further examine the differences. p<0.05, LPS alone vs. vehicle control; p<=NS, B+LPS vs. vehicle control; p=NS, FP+LPS vs. vehicle control; p<0.05, LPS alone vs. B+LPS; p<0.02, LPS alone vs. FP+LPS; p<0.01, B+LPS vs FP+LPS.

#### Wet:Dry ratio

Lung wet to dry ratios were measured in separate treated mice. These treated animals were not subjected to lung lavage. In vehicle control treated mice, the weght of the lung was 3.5 ± 0.4 g and increased to 6.8 ± 1.2 g following LPS treatment (p<0.05). The lung W/D ratio decreased to 5.2 ± 1.1 g for mice receiving Bignay but was not statistically different from LPS treated mice (p=NS). However, the blockade caused by FP was near the baseline value (p<0.05 vs LPS alone).

#### Airway inflammatory cell lung infiltration

LPS caused inflammatory cell infiltration in the lung interstitium, surrounding the airways and pulmonary blood vessels (Fig 5B) compared to vehicle control treated mice (Fig 5A). A marked reduction of migrated cells to the site of inflammation caused by LPS stimulation was reduced by 1000 mg.kg^-1^ Bignay (Fig 5C). Bignay was as effective as FP in reduction of cell infiltrate (Fig 5D) suggesting that Bignay could be a good candidate in protecting the migration of inflammatory cells to the site of inflammation.

**Fig 5.**
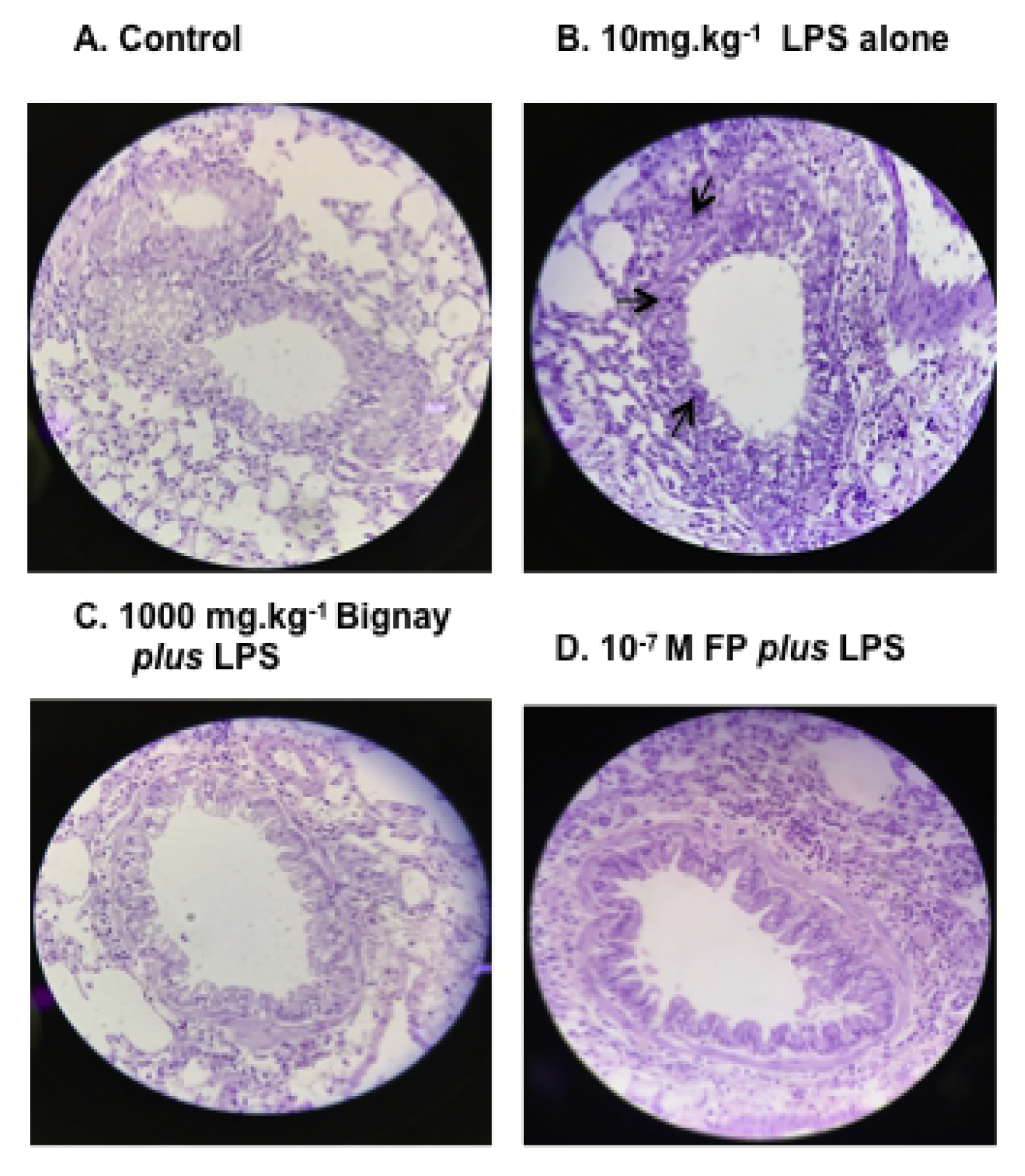
Infiltration of inflammatory cells after instillation of LPS. Representative microsectioned of airways and lung parenchyma for histologic assessment of cellular infiltration of inflammatory cells. Airway inflammation was analyzed after staining the lung with hematoxylin and eosin dye. (A). Vehicle control showed normal appearance of the airway and lung parenchyma with minimal inflammatory cells in lung tissue (B). LPS-induced ALI caused accumulation of inflammatory cells surrounding the airways and within epithelium (see black arrow). (C). Treatment with 1000 mg.kg^-1^ *Antidesma bunius* (L.) Spreng [Bignay] caused reduction of migrated inflammatory cells in the airway that is comparable with vehicle control mice (control). (D). Administration of Fluticasone Propionate (FP) also caused reduction of cell migration caused by LPS stimulation. Light microscopic magnification was set at X60 pixel.

#### Pulmonary Vascular Leak

Lastly, we examined the effect of Bignay fruit extracts on the pulmonary vascular permeability caused by LPS. Injection of Evans Blue dye [46] into the tail vein caused severe blue coloration of the lungs 2 hrs after administration of LPS, signaling the induction of vascular leakage. In the vehicle control-treated mice, the appearnce of lung remained pinkish in color (Fig 6A), but in LPS-induced acute lung injury, blue coloration of the lung was evident 2 hrs later (Fig 6B). In animals with 500 mg.kg-1 Bignay extract, there was a minimal blockade of vascular leak (Fig 6C). Increasing the dose of Bignay to 1000 mg.kg^-1^ substantially blocked the vascular leakage caused by LPS (Fig 6D); Bignay was as effective as FP (Fig 6E). These data provide preparatory evidence for Bignay’s role, like FP, in blocking the LPS-induced pulmonary vascular leak.

**Fig 6.**
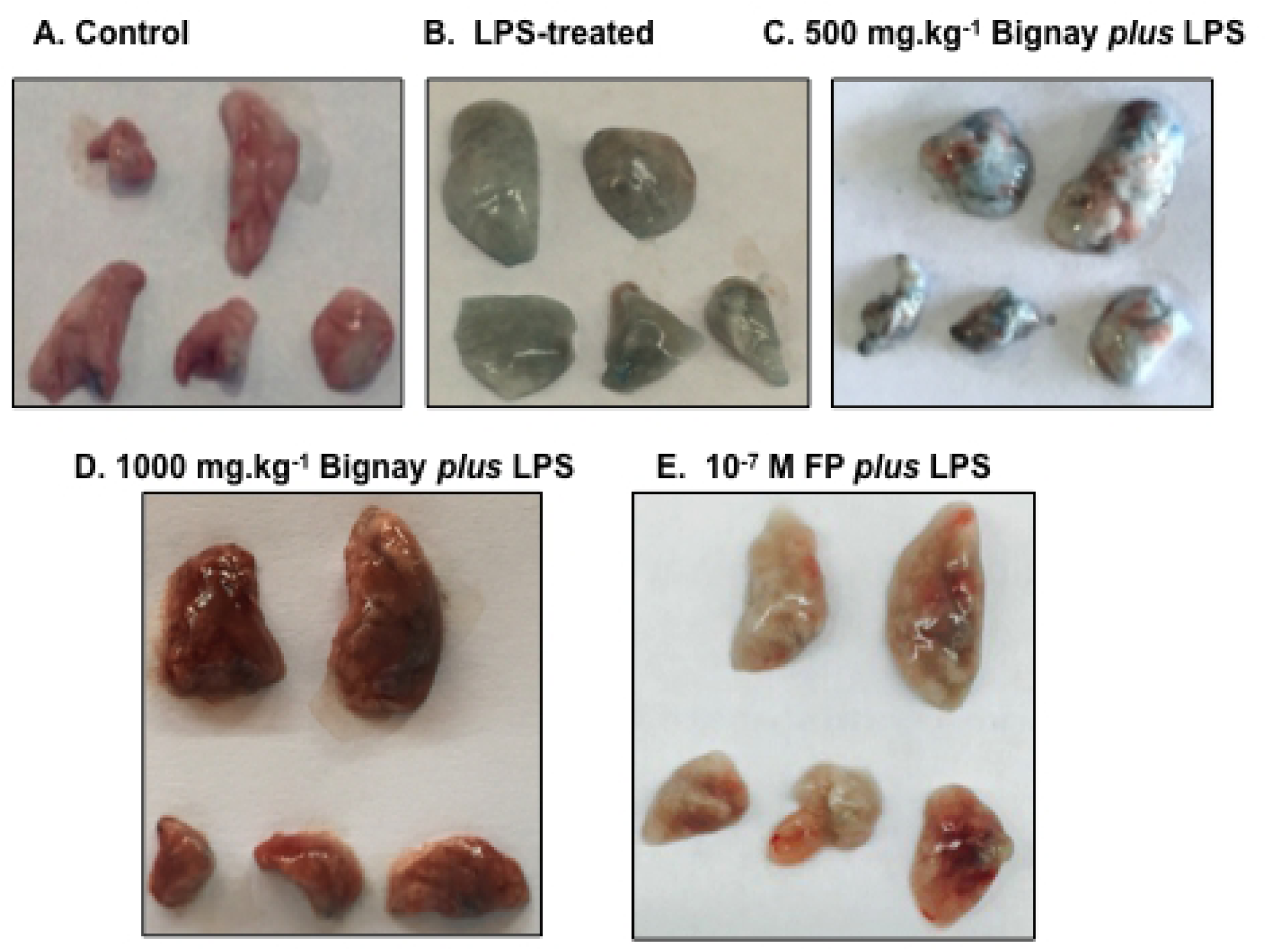
Assessment of Lung Vascular Leak. Evans Blue Dye is a known dye for assessing lung damage caused by LPS in mice. (A) vehicle control treated mice; (B) The blue coloration of lung indicates the vascular leak by LPS. Pretreatment of mice with 500 mg.kg^-1^ Bignay (C) caused minimal effect in preventing vascular leak, however, administration of 1000 mg.kg^-1^ Bignay (D) via *i.p.* injection substantially prevented the vascular leak caused by LPS stimulation. The positive reference drug (E), Fluticasone Propionate (FP), protected the effect of LPS in induction of acute lung injury. The effect of *Antidesma bunius* (L.) Spreng [Bignay] is as effective as FP.

## Discussion

Philippine berries play an important role in inflammatory disease control [9,10,21–24,26]. The objective of this study is to investigate the protective and toxicological effects of the ethanolic extract of *Antidesma bunius* (L.) Spreng [Bignay] fruit (Fig 1) on biomarkers of ALI caused by LPS and compared with Fluticasone Propionate, a synthetic steroid of the glucocorticoid drug family.

We first analyzed the amount of heavy metals (Table 1) as well as total aflatoxins and aflatoxins B1 (Table 2) to ensure that unwanted substances are not present in the Bignay fruit extracts, According to the Philippine Accredited Institute of Pure and Applied Chemistry, Bignay samples have insignificant trace of heavy metals. Another findings showed that in the Guidelines on the Registration of Herbal Medicines of the Department of Health, total aflatoxins and aflatoxins B1 were below the limit stipulated. The samples used in these analyses are then recommended to provide evidence that Bignay is safe (Table 1 and Table 2) for use in laboratory animals and likely, in humans.

**Table 2.** Analysis of Total Aflatoxin and Aflatoxin B1 (n=3 trials)

| Trial Number | Total Aflatoxin (ppb) | Aflatoxin B1 (ppb) |
| --- | --- | --- |
| 1 | 13.26 | 4.27 |
| 2 | 12.33 | 5.16 |
| 3 | 12.64 | 4.49 |
| Average | 12.75 ± 0.47 | 4.64 ± 0.46 |

The flavonoid, phenolic, and steroid contents of ethanolic Bignay fruit extracts were next quantified. Flavonoids exemplify the most widely distributed group of plant phenolics and are abundant in foods [27, 28]. It has been reported that flavonoids are particularly beneficial, acting as antioxidants and providing protection against various inflammatory diseases [27,28,48,49]. While phenolics account for the majority of antioxidant activity in plants or plant products, they are the largest group of phytochemicals [28,38,48,50]. In patients with cardiovascular disease, a type of inflammatory disease, plant sterols have the potential to block dietary cholesterol absorption. With these results, it could be postulated that Bignay fruits could play an important role in radical scavenging and can therefore be considered as beneficial plant species for natural antioxidant sources with potential use for the treatment of many life-threatening inflammatory diseases.

It is likely that the inhibition of LPS-induced ALI could be attributed to the active secondary metabolites, phenols, flavonoids, and steroids, measured in Bignay fruit extracts. Nevertheless, the different functions of these metabolites are not well understood yet. The effects of natural herbal compounds in inflammatory cell infiltration have previously been demonstrated in the murine asthma model [51, 52]. The *Boerhavia procumbens* methanolic extract showed a decrease in the number of eosinophils and T-cells in toluene diisocyanate lung exposed rats [52]. *Octimum gratisimum*, a widely used folk medicine in Brazil, induced a decrease in airway inflammation and development of cytokines in allergic mice [51]. In addition, treatment with flavonoid quercitin in immunized mice also reduced the number of inflammatory cells and lung homogenate cytokine levels [50].

Using the murine model of LPS-induced ALI, we demonstrated that administration of LPS increased: a) lung inflammatory cell number (Fig 2), b) total lung protein content (Fig 3), c) wet to dry ratio, d) endogenously secreted IL-6 from BALF (Fig 4), e) airway inflammation (Fig 5) and f) pulmonary vascular leak (Fig 6). Administration of Bignay fruit extracts and FP was apparently, equally effective in preventing the LPS-induced ALI (Figs 2,3,4,5) and pulmonary vascular leakage (Fig 6). The use of ANOVA with Tukey test was used to test for population mean differences; mean of all animals treated with Bignay, mean of all animals treated with FP, which is considered to be the best available method in cases where confidence intervals are required. We noticed that there was no statistical significance when Bignay+LPS was compared to FP+LPS [Figures 3-4]; however, the protective effect of FP differed from Bignay in the number of total lung cell [Fig 2], suggesting that FP is more efficacious than Bignay in preventing airway inflammation.

LPS administration increased the number of inflammatory cells migrating into the lungs (Fig 2) and the overall BALF protein content of the lungs (Fig 3). In blocking these responses, the protective effect of extracted Bignay with a high amount of quercitin (flavonoid content) in the extracts is comparable to FP. In addition, it is evident that Bignay’s anti-inflammatory properties are linked to the combination of phenols, steroids and flavonoids, which are considered to have anti-oxidant and anti-inflammatory properties. Previous studies have shown that components of flavonoids and phenolic acids play a significant role in cancer and other human inflammatory disease regulation [48, 53].

Airway pro-inflammatory mediators such as secreted cytokines have been well known to lead to persistent chronic airway inflammation and airway remodeling [4,5,7,8]. Previous studies have shown that FP has significantly suppressed secreted interleukins [42,43–45]. Here, the protective effect of the natural Bignay fruit extract is as effective as FP in inhibiting the release of IL-6 in animals receiving LPS (p<0.05 vs LPS alone; Fig 4), indicating that Bignay could also play a significant modulatory role in preventing the remodeling phase of the airway as IL-6 is one of the ALI mediators.

The lung wet-to-dry (W/D) weight ratio was used as an index of lung water accumulation after the instillation of LPS. As a result, the rise in lung edema following LPS instillation was a combination of an inflammatory response and a disruption in fluid exchange, both of which are caused by extracellular LPS’s direct action. Unlike FP, the administration of Bignay, did not differ significantly in blocking the formation of lung edema but may presumably be sufficient to block lung edema caused by LPS in longer period of treatment.

As observed in essential structural changes in lung morphology, LPS caused lung damage (Fig 5). Increases in airway smooth muscle thickness (hypertrophy) and epithelial cell denudation were evident in H&E stained tissues. However, after treatment with either Bignay fruit extract or FP, the alteration in lung morphology returned to near normal structural appearance; this is an indication that Bignay fruit extract could prevent the process of airway remodeling in murine model of ALI. Herbal formula has been shown to reduce cell infiltration in the lungs and secretion of TNFα and IL-6 in an emphysema-induced model of elastase plus LPS [54]. In this regards, previous studies have shown that not only can herbal medicine suppressed BALF inflammatory cells, but also cytokine levels and airway remodeling. The influence of the traditional medicinal product *Callicarpa japonica* on the prevention of inflammatory diseases reduced neutrophil influx and IL-6 production in the cigarette smoke model (COPD) supported these findings [55–57].

ALI/ARDS animal models have been used as features to help understand the pathogenesis of this lung disease, in particular the evaluation of vascular permeability after lung damage induction. In mice acutely exposed to LPS, Dudek *et al*. showed that neutralizing mAb treatment directed against Group V Phospholipase A_2_ (57) reduced inflammatory activities of the airway mediated in part by neutrophilic infiltration with alteration of F-actin assembly and gap junction, VE-Cadherin [36]. To date, no studies have tested the preventive effects of medicinal plants in murine model of LPS-induced ALI/ARDS *in vivo*. This research is the first demonstration to compare the anti-inflammatory efficacy of Bignay fruit extract with Fluticasone Propionate (FP) in blocking ALI induced by LPS (Fig 7).

**Fig 7.**
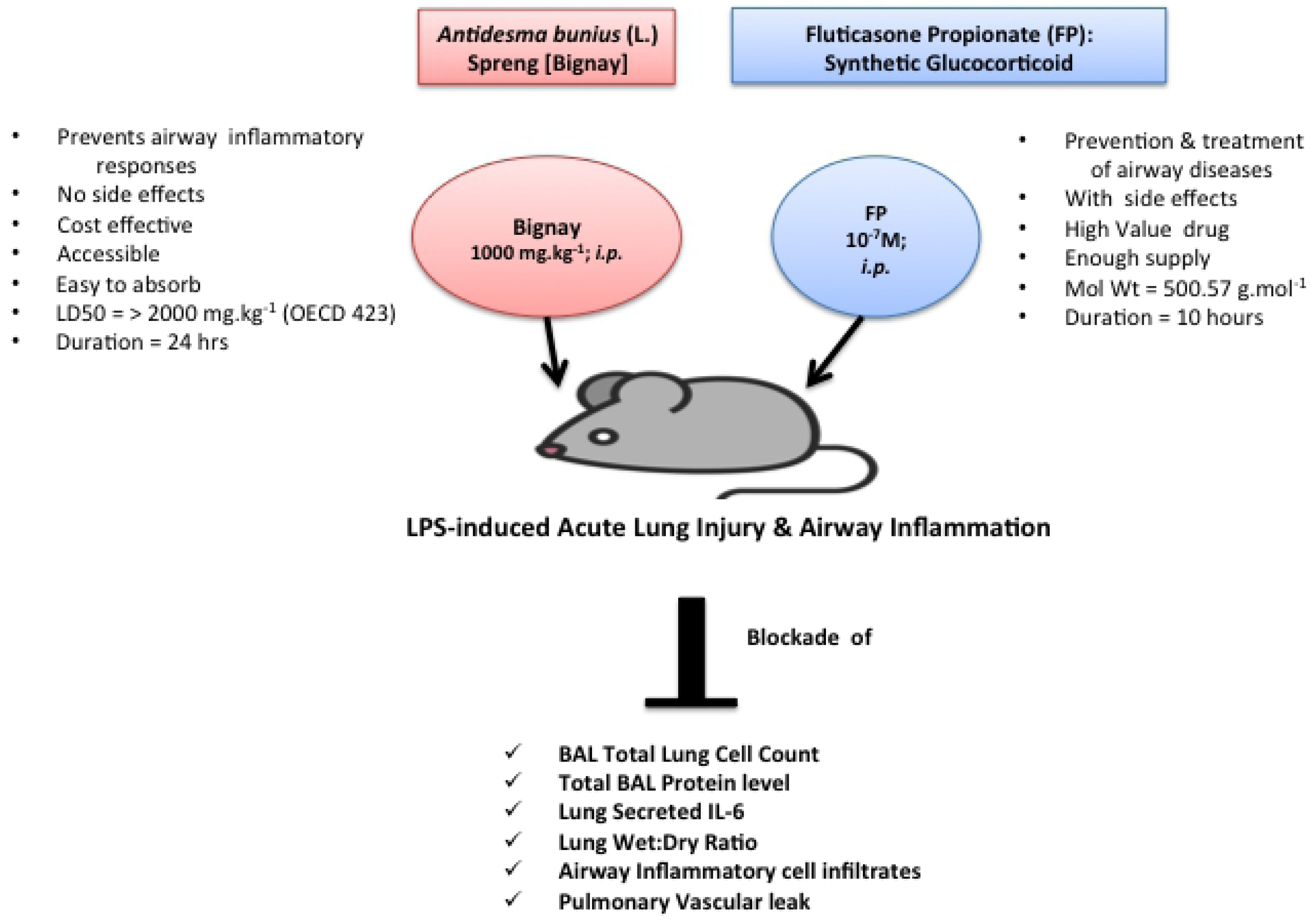

The beneficial properties of Bignay have been anecdotally recognized and have developed a powerful suppressive effect on inflammatory diseases, whereas FP belongs to a class of drugs known as corticosteroids and is used for the long-term treatment of upper and lower airway diseases such as asthma and COPD [42–45]. In this study, both Bignay and FP have been shown to be similarly successful and well tolerated by LPS-treated mice (Fig. 2-4), indicating that Bignay fruit extract, free of pollutants and heavy metals, could be converted into a healthy, clinically based herbal drug. In the end, Bignay would be able to increase local livelihood of farmers and would provide a fair, protected and reliable herbal anti-inflammatory medicine for a variety of inflammatory diseases.

It is important to note some limitations of our findings. Although our findings showed preventive and non-toxic effects of Bignay fruit extracts in LPS-induced ALI in *vivo*, our results are focused on an ALI murine model. At least to our knowledge, we have evaluated the effects of Bignay extracts in the murine model of ALI, but not all ALI indices have been evaluated. However, the mechanisms involved in inhibiting ALI triggered by LPS stimulation have been partly established. In addition, the isolation or structural formula of active compounds in Bignay was not included in this analysis, but biochemical compound recognition plus the anti-inflammatory activity of the three secondary metabolites were quantified. Although the content of total aflatoxins and aflatoxin B1 were determined, the source of *Aspergillus Flavus* and *Apergillus parasiticus* were not identified in this study. Histological airway analysis is close to that of humans with ALI; our data in mice, however, cannot be explicitly extrapolated to the human state.

As for ALI in humans, therapy does not confer therapeutic effectiveness after significant histological changes occur. Although the blocking effects on ALI production in this study are relatively short, a hyper-acute model of ALI is the model used here, eliciting a rapid response after LPS infusion. By contrast, in humans, ALI typically grows over a considerably longer period (hrs to days), as in a cumulative rather than square wave fashion, the inciting stimulus likely causes lung injury. Therefore, the demonstration that Bignay administered pre-LPS is successful in abolishing or altering ALI in mice for up to 2 hrs indicates the likelihood of a longer window for intervention in the more chronic human production of ALI. This depends, of course, on the degree to which the results in this study translate into the human situation. It is also presumed that until ALI is fully manifested, this therapy is probably helpful.

We also observed that complete blockade of ALI caused by Bignay is not all done in treated mice. Nonetheless, with 1000 mg.kg-1 Bignay, a significant or near complete reduction of the impact of LPS on ALI can be induced, especially in the reduction of vascular leakage caused by LPS stimulation (Fig 6). This blocking effect is as efficient as FP in LPS-induced ALI/ARDS in mice, indicating that Bignay is highly likely, could prevent the development of ALI/vascular leakage caused by bacterial soluble factor such as LPS.

## Conclusion

Our research is the first demonstration that the inhibition by *Antidesma bunius* (L.) Spreng fruit extracts of LPS-induced ALI is as effective in mice *in vivo* as Fluticasone Propionate (FP), a corticosteroid. Figure 7 represents the schematic diagram comparing Bignay’s and FP’s physiognomies and preventive effects on LPS-induced ALI. Treatment with either Bignay or FP resulted in a substantial reduction in the number of lung inflammatory cells, lung protein content, lung edema, and vascular pulmonary leakage caused by LPS. Most importantly, we demonstrated that secreted IL-6, one of the biomarkers of ALI, was inhibited by Bignay fruit extract; Bignay’s inhibitory effect on secreted IL-6 was only ∼55%, but comparable to FP.

In accordance with these findings, in the presence of Bignay or FP in treated mice, a substantial reduction in vascular leakage was achieved. The precise mechanism by which Bignay modulates ALI, unlike in previous FP studies, is unknown, but it is possible that Bignay’s ability to attenuate ALI and vascular leak indices can be explained, at least in part, by our new findings. The quantified phenols, phytosterols and flavonoids showed contributory or synergistic inhibitory effects on LPS-induced ALI biomarkers. Collectively, Bignay may provide a therapeutic strategy for human ALI regulation that may be useful for new drug discovery with reduced side effects in inflammatory conditions. The prevention of ALI in humans by Bignay remains further investigations, e.g., clinical trials.

## Acknowledgments

We are grateful for the access of the animal laboratory facility to the College of Veterinary Medicine, Cagayan State University and R&B Facility, St. Luke’s Medical Center, Philippines. The writers are thankful for the support and critical analysis of the manuscript by Professor Urdujah Gaerlan Alvarado, Professor Kozo Watanabe, Dr. Brigido L. Carandang Jr., Dr. Swapan Chatterjee and Dr. Charina de Silva. Authors would also like to acknowledge the assistance provided in the H&E staining of the lung tissue parts by the Department of Histopathology at Cagayan State University; we are grateful to JL, KSHB, RCD and JILR, who during the research time took care of the mice and their assistance in conducting the experiments.

## Author Contributions

Conceptualization: Maria Nilda M. Muñoz, Kozo Watanabe Data curation: Maria Nilda M. Muñoz, Kozo Watanabe, Swapan Chatterjee Funding acquisition: Maria Nilda M. Muñoz, Jennifer Lucero, Urdujah Gaerlan Alvarado,

Investigation: Maria Nilda M. Muñoz, Urdujah Gaerlan Alvarado, Kozo Watanabe

Methodology: Maria Nilda M. Muñoz, Charina de Silva, Jennifer Lucero, Kimberly Stacy Hope Benzon, Reuel C. Delicana, Jerica Isabel L. Reyes

Project administration: Maria Nilda M. Muñoz, Urdujah Gaerlan Alvarado, Kozo Watanabe

Writing – original draft: Maria Nilda M. Muñoz

Writing – review & editing: Maria Nilda M. Muñoz, Kozo Watanabe, Swapan Chatterjee

